# Networks of HIV-1 envelope glycans maintain antibody epitopes in the face of glycan additions and deletions

**DOI:** 10.1101/2020.02.21.959981

**Authors:** Gemma E. Seabright, Christopher A. Cottrell, Marit J. van Gils, Alessio D’addabbo, David J. Harvey, Anna-Janina Behrens, Joel D. Allen, Yasunori Watanabe, Allison Maker, Snezana Vasiljevic, Natalia de Val, Rogier W. Sanders, Andrew B. Ward, Max Crispin

## Abstract

Numerous broadly neutralizing antibodies (bnAbs) have been identified that target the glycans of the HIV-1 envelope spike. Neutralization breadth is notable given that glycan processing can be substantially influenced by the presence or absence of neighboring glycans. Here, using a stabilized recombinant envelope trimer, we investigate the degree to which mutations in the glycan network surrounding an epitope impact the fine glycan processing of antibody targets. Using cryo-electron microscopy and site-specific glycan analysis, we reveal the hierarchy of importance of glycans in the formation of the 2G12 bnAb epitope, and show that the epitope is only subtly impacted by variations in the glycan network. In contrast, we show that the PG9 and PG16 glycan-based epitopes at the trimer apex are dependent on the presence of the highly conserved surrounding glycans. Glycan networks underpin the conservation of bnAb epitopes and are an important parameter in immunogen design.

## INTRODUCTION

The envelope spike (Env) of the human immunodeficiency virus type 1 (HIV-1) mediates infection of target host cells and is consequently a main target for vaccine design. However, Env displays extreme antigenic diversity, meaning only an immune response of exceptional breadth will be protective (Burton et al., 2012). Additionally, a dense coat of host derived, immunologically ‘self’ N-linked glycans shield the underlying protein from host antibody responses (Wei et al., 2003). In spite of these hurdles, approximately a third of infected individuals develop broadly neutralizing antibody (bnAb) responses against Env after several years of infection (Simek et al., 2009; van Gils et al., 2009).

While the Env glycan shield typically limits antibody neutralization, many bnAbs have, paradoxically, evolved to recognize epitopes that are either entirely or partially composed of N-linked glycans (Blattner et al., 2014; Doores and Burton, 2010; Falkowska et al., 2014; Huang et al., 2014; McLellan et al., 2011; Pancera et al., 2013; Pejchal et al., 2011; Scharf et al., 2014; Walker et al., 2011; Walker et al., 2009). These bnAbs recognize the glycans at four distinct regions of Env: the gp120/gp41 protomer interface (e.g. PGT151), surrounding the CD4 binding site (e.g. HJ16), the V1/V2 loops at the trimer apex (e.g. PG9 and PG16), and the oligomannose-type glycans centered around the highly conserved N332 site on the outer domain of gp120 (e.g. PGT135 and 2G12) (Crispin et al., 2018). Given the number of bnAbs targeting the N332 glycan, it has previously been termed the ‘supersite of immune vulnerability’ (Kong et al., 2013).

It is well established that the passive transfer of bnAbs protects non-human primates and humanized mice from viral challenge (Pegu et al., 2017; van Gils and Sanders, 2014). Thus, bnAbs are now being investigated for both therapeutic use (Stephenson and Barouch, 2016), and to guide the design of Env-based immunogens intended to elicit similarly broad and neutralizing responses (Burton, 2017; Sanders and Moore, 2017). The latter approach typically involves producing recombinant mimics of the native, virion-associated Env trimer that present multiple bnAb epitopes, and/or immunogens specifically designed to target the germline-encoded bnAb precursors (gl-bnAbs) (Sanders and Moore, 2017; Stamatatos et al., 2017).

Currently, the most widely studied recombinant Env mimics are the BG505 SOSIP.664 trimers, based on the subtype A transmitted/founder virus sequence, BG505. Various modifications, including the introduction of a disulfide bond (SOS), an isoleucine to proline mutation (IP), and truncation at the 664 position (.664), increase both the stability and solubility of the trimers (Sanders et al., 2013). The resulting trimers display native-like structure and antigenicity (Sanders et al., 2013; Ward and Wilson, 2017), and are lead candidates in ongoing human immunogenicity studies (Dey et al., 2018; NCT03699241).

The BG505 transmitted/founder virus naturally lacks the conserved N332 glycan site, thus this glycan was also included (T332N) in the BG505 SOSIP.664 trimers in order to introduce the ‘supersite’ epitope (Sanders et al., 2013). However, the BG505 sequence also lacks glycans at the 241 and 289 positions, despite their presence in 97% and 72% of HIV-1 isolates, respectively. The presence or absence of holes within the glycan shield has recently received a lot of attention because of the putative role of holes in initiating neutralizing antibody (nAb) responses.

Immunization of rabbits with BG505 SOSIP.664 trimers elicits autologous nAb responses centered on glycan holes at positions 241 and 289 (Klasse et al., 2016; McCoy et al., 2016). Filling this hole (i.e. by introducing a glycan site) blocks antibody neutralization (McCoy et al., 2016). Similar results have been observed with nAbs targeting holes at the 130, 197 and 465 positions in immunogenicity studies with native-like trimers from different isolates (Crooks et al., 2017; Crooks et al., 2015; Klasse et al., 2018; Klasse et al., 2016; Voss et al., 2017). Such autologous nAb responses can be readily redirected *in vivo* by closing the glycan holes, and opening new ones elsewhere on the trimer (Ringe et al., 2019). This phenomenon is echoed in natural infection, as the glycan shield ‘shifts’ to escape arising nAbs (Dacheux et al., 2004; Moore et al., 2012; Wagh et al., 2018; Wei et al., 2003). The N332 glycan, for example, has been observed to shift from the N334 position and back again after the appearance of nAbs (Moore et al., 2012). While it is accepted that glycan holes offer an immunodominant distraction capable of eliciting autologous nAbs, the extent to which holes hinder the development of bnAbs remains largely unknown. There is evidence to suggest that more complete glycan shields in transmitted/founder viruses correlate with the development of greater neutralization breadth in infected individuals (Wagh et al., 2018). Future immunization strategies may therefore include immunogens with closed glycan holes, in order to redirect the nAb response away from the immunodominant protein surface toward more broadly neutralizing glycan-based epitopes (McCoy et al., 2016; Ringe et al., 2019).

The elicitation of a bnAb response requires the activation of bnAb precursor B cells. Effective immunogens must therefore be capable of engaging the B cell receptor (i.e. the gl-bnAb), prior to affinity maturation of the bnAb in the germinal centers. However, this process is hampered by the low affinity of gl-bnAbs to Env, often due to their inability to accommodate conserved N-linked glycans (Doores et al., 2013; Hoot et al., 2013; Ma et al., 2011; McGuire et al., 2014; Xiao et al., 2009). Thus an alternative, albeit closely linked, approach to eliciting bnAbs, is to prime with glycan-depleted immunogens, capable of engaging gl-bnAbs, and subsequently boost with their ‘filled-in’ derivatives in order to drive the development of neutralization breadth (Jardine et al., 2013; McGuire et al., 2013; Medina-Ramirez et al., 2017; Stamatatos et al., 2017; Steichen et al., 2016).

Glycan density, however, impacts glycosylation processing, which can in turn influence epitope presentation. The unusually high density of N-linked glycans on gp120 limit the extent to which individual sites can be processed by the host’s α-mannosidases (Behrens and Crispin, 2017). Thus, gp120 displays a significant population of under-processed oligomannose-type glycans, termed the intrinsic mannose patch (IMP) (Bonomelli et al., 2011; Doores et al., 2010a; Go et al., 2013; Pritchard et al., 2015a). Analysis of recombinant, monomeric gp120 revealed that the removal of individual glycan sites from within the IMP often results in larger-than-expected decreases in the abundance of oligomannose-type glycans, as sites surrounding the deletion become more susceptible to glycan processing (Pritchard et al., 2015a). In Env trimers displaying native-like conformations, additional steric hindrances imposed by glycan and protein elements from neighboring protomers give rise to a further trimer associated mannose patch (TAMP) (Behrens et al., 2017a; Cao et al., 2017; Pritchard et al., 2015c). Analysis of glycan-depleted, trimeric immunogens also revealed increased glycan processing at sites proximal to the glycan deletions (Behrens et al., 2018; Cao et al., 2017). Furthermore, a number of studies have reported correlations between glycan density and the abundance of under-processed oligomannose-type glycans (Coss et al., 2016; Stewart-Jones et al., 2016). Thus, while oligomannose-type glycans are a conserved feature of the Env glycan shield, and a key bnAb target, in some circumstances they can become susceptible to enzymatic processing.

Given the propensity for glycan density to influence the processing of glycans, we sought to determine the impact of individual glycan site additions and deletions on bnAb epitopes. Here, using glycopeptide analysis of BG505 SOSIP.664 trimers, we reveal that glycan site addition and deletion influences the fine processing of glycans both proximal to the mutated glycan site, and elsewhere on the trimer. We further probe the tolerance of bnAbs to glycan mutations, and reveal the differing dependencies of mannose patch-targeting and apex-targeting bnAbs on the surrounding N-linked glycan sites. We also report a high-resolution structure of the 2G12 bnAb in complex with the BG505 SOSIP.664 trimer by cryo-electron microscopy (EM) and reveal details of the wider network of glycans that maintain the epitope. Furthermore, we show the N334 to N332 escape mutation minimally impacts glycosylation processing. The diverse impact of glycan holes on glycan-dependent bnAbs underscores the role of glycopeptide analysis in vaccine design and the development of new immunogens.

## RESULTS

### Enhanced mapping of the 2G12 epitope by cryo-EM

To elucidate the molecular details of the 2G12 epitope, we solved a 3.8 Å structure of 2G12 Fab_2_ bound to BG505 SOSIP.664 by cryo-EM (**Figure 1A, Figure S1, Table S1**). The two Fabs of the 2G12 bnAb are known to adopt a domain-swapped dimer conformation (Fab_2_) via exchange of their VH domains (Calarese et al., 2003; Doores et al., 2010b). This unique architecture creates two primary binding sites at the VH/VL interfaces and two secondary binding sites on either side of the VH/VH interface. The 2G12 Fab_2_ binds to a 1,879 Å^2^ epitope composed entirely of N-linked glycans (Calarese et al., 2005; Sanders et al., 2002; Scanlan et al., 2002; Trkola et al., 1996). The primary binding sites make contact with the terminal α 1,2-linked mannose residues of the D1-arms of the oligomannose-type glycans at positions N392 and N295 (**Figure 1A-C**; see **Figure S5** for nomenclature). The secondary binding sites make contact with the D2- and D3-arms of the oligomannose-type glycans at positions N332 and N339 (**Figure 1A-C**). Although not directly in contact with the 2G12 Fab_2_, the glycans at N363 and N386 may also play a role in 2G12 binding by providing support to the N392 glycan via glycan/glycan interactions (**Figure S1B**) (Sanders et al., 2008). Previous studies have reported a potential contact between the 2G12 Fab_2_ and glycans at positions N137 and N411 (Chuang et al., 2019; Murin et al., 2014), however, no coordinated density was observed for these glycans in our reconstruction, indicating they do not play a direct structural role in 2G12 binding (**Figure S1C**).

**Figure 1.**
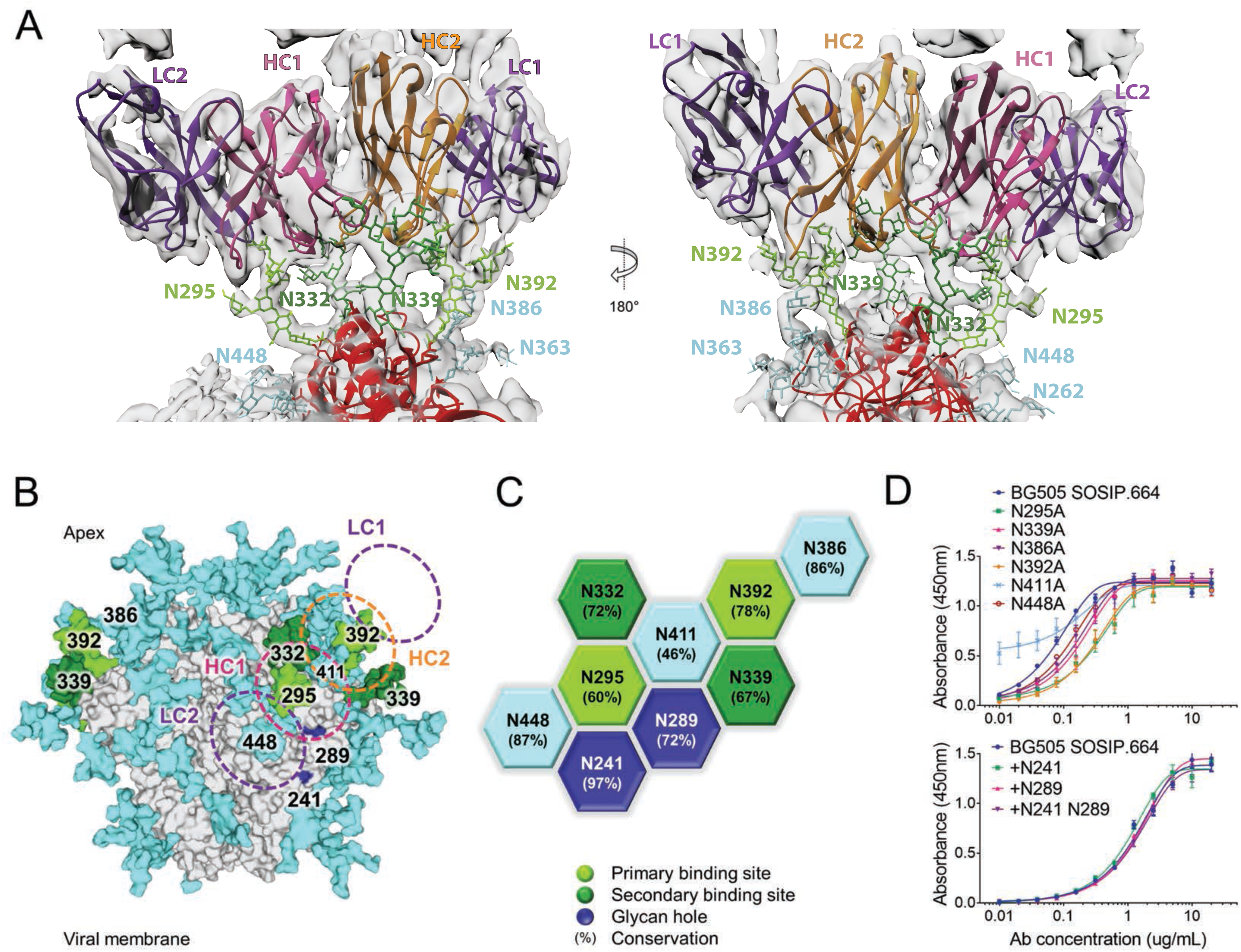
The 2G12 epitope. A) The cryo-EM structure of the 2G12 Fab_2_ in complex with BG505 SOSIP.664 was resolved to 3.8 Å. Electron density is shown in grey. No density was observed for the glycan at N411 (**Figure S1**). Glycans bound by the 2G12 primary binding site are colored light green, glycans bound by the secondary binding site are colored dark green, the surrounding network of glycans are shown in cyan. B) Model of a fully glycosylated BG505 SOSIP.664 trimer based on PDB 5ACO with glycans added according to Behrens et al., 2016 (Behrens et al., 2016). The glycan holes at the 241 and 289 positions are highlighted in dark blue. The footprint of 2G12 is shown for orientation. C) The network of glycans surrounding the 2G12 epitope, and their conservation (**Figure S2**). D) 2G12 binding to glycan knockouts and knock-ins was assessed by ELISA, mean ±SEM.

To further gauge the contribution of individual glycans to the 2G12 epitope, we assessed antibody binding to glycan knockouts by ELISA. As expected, the N295A and N392A deletions resulted in the largest decrease in antibody binding (**Figure 1D**). The N448A and N386A knockouts, while not directly contributing to the 2G12 epitope, diminished binding to a similar extent as a secondary binding site deletion. In contrast, the N411A mutation increased 2G12 binding. Additionally, we sought to determine the impact of filling the nearby 241/289 glycan hole on 2G12 binding. Neither the +N241 or +N289 glycan knock-ins, nor the +N241 N289 double knock-in had a substantial impact on 2G12 binding (**Figure 1C**).

### Overall resilience of the mannose patches to glycan addition and deletion

To assess whether the observed differences in 2G12 binding could be attributed to differences in glycan processing, the glycosylation profiles of the glycan site mutants were compared to that of BG505 SOSIP.664 trimers (**Figure 2**). As the removal of glycan sites can conceivably negatively impact trimer integrity, a quaternary structure-dependent antibody (PGT145), targeting a region distal to the site of mutation, was used for purification. The N-glycans from the target glycoproteins were enzymatically released, fluorescently labeled, and analyzed by hydrophilic interaction liquid chromatography-ultra performance liquid chromatography (HILIC-UPLC; **Figure S3**). Quantification of oligomannose-type species was performed by the integration of chromatograms before and after digestion with Endoglycosidase H (Endo H; **Figure 2B**). Additionally, the oligomannose-type glycans from three biological replicates of BG505 SOSIP.664 were quantified (**Figure 2C**), revealing a coefficient of variation of 2.2% for Man_9_GlcNAc_2_ and 1.5% for total oligomannose-type glycans.

**Figure 2.**
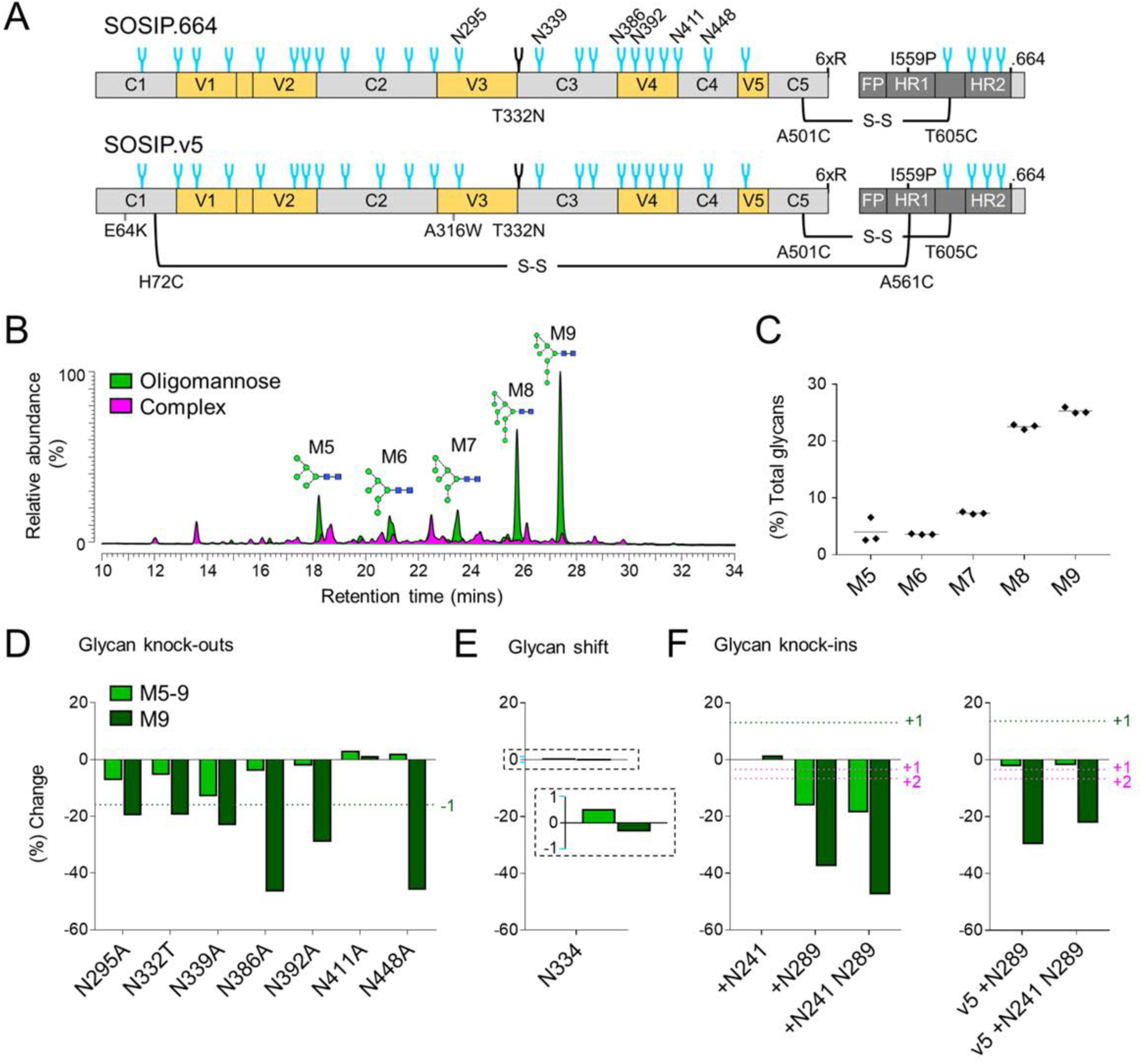
Impact of glycan site additions and deletions on overall glycosylation profiles. A) Linear schematic of the BG505 SOSIP.664 and SOSIP.v5 trimers, with stabilizing mutations annotated. B) Example HILIC-UPLC profile of fluorescently labeled N-linked glycans released from BG505 SOSIP.664. Oligomannose-type glycans (green) are quantified by integration of peaks before and after digestion with Endo H (pink). C) Quantification of individual oligomannose-type glycans from three biological replicates of BG505 SOSIP.664. D-F) Effect of glycan site deletion, shift, or addition on the abundance of Man9 (M9) and total oligomannose-type glycans (M5-9). Values represent the percentage change in abundance, relative to BG505 SOSIP.664: percentage change = [(% glycan mutant - % BG505 SOSIP.664)/% BG505 SOSIP.664] × 100. The dashed green lines represent the change in Man9 expected upon either the deletion (D) or addition (F) of a glycan site comprising solely Man9 structures. The dashed magenta line (F) represents the decrease in the abundance of Man9 expected upon the addition of one or two sites comprising solely complex-type glycans.

As per BG505 SOSIP.664 trimers, the chromatograms of all the glycan mutants were dominated by oligomannose-type glycans, though the distribution of individual oligomannose-type glycans, particularly Man_9_GlcNAc_2_ and Man_8_GlcNAc_2_ varied slightly (here referred to as Man9, Man8 etc.) (**Figure S3**). Previous site-specific glycan analyses of BG505 SOSIP.664 trimers have reported the N295, N332, N339, N386, N392, and N448 sites to be occupied by oligomannose-type glycans, predominantly Man9 (Behrens et al., 2016; Cao et al., 2017). Accordingly, deletion of each of these sites resulted in a decrease in the abundance of both total oligomannose-type glycans and Man9 structures (**Figure 2D**). The largest changes were observed for the N448A and N386A glycan-knockouts, which resulted in a 46% and 47% decrease in the abundance of Man9, respectively. Many of the observed decreases were somewhat larger than the decrease predicted upon the loss of a glycan site comprising solely Man9 structure (16%; **Figure 2B**, dashed line). Thus, glycan site deletion on BG505 SOSIP.664 trimers can result in widespread increased glycosylation processing. This effect was not universally observed, for example, the N411A glycan site knockout had minimal impact on overall glycosylation processing despite its location at the center of the IMP (**Figure 1 and 2D**).

Additionally, we investigated the impact of the N332 to N334 glycan ‘shift’ escape mutation on glycosylation processing. In contrast to the deletion of the N332 site, the migration of the glycan to the N334 position had negligible impact on either the abundance of Man9, or total oligomannose-type glycans (**Figure 2E**).

Given that glycan site deletion generally resulted in increased glycosylation processing, and a glycan shift mutation did not impact glycosylation processing, we hypothesized that glycan site addition may restrict processing. Analysis of the +N241 glycan knock-in, however, revealed only a minimal increase in the relative abundance of Man9 structures (2%; **Figure 2F**). In contrast, the +N289 and +N241 N289 knock-ins resulted in a decrease in both the abundance of oligomannose-type glycans, and Man9 structures. This result may be expected if the knocked-in sites were composed of predominantly complex-type glycans. However, the observed decrease in the abundance of Man9 (38% and 48%, respectively) far exceeds the predicted decreases upon the addition of one or two sites containing only complex-type glycans (3% and 7%, respectively; **Figure 2F**, dashed line). Thus, both glycan site deletions and additions including the N289 site appear to be increasing the glycosylation processing of the trimer.

We hypothesized that the BG505 SOSIP.664 trimer may be unaccommodating of the N289 glycan, and that its addition may be inducing conformational changes which in turn influence glycosylation processing. We therefore repeated the analysis of the +N289 and +N241 N289 knock-ins on hyperstabilized BG505 SOSIP.v5 trimers (**Figure 2A**), which incorporate further stabilizing mutations, including an additional inter-subunit disulfide bond, and display reduced conformational flexibility (Torrents de la Pena et al., 2017). While the decrease in the abundance of Man9 on the SOSIP.v5 background was not as severe as on the SOSIP.664 trimers, it still exceeded the decrease predicted if the knocked-in sites were composed of only complex-type glycans (**Figure 2F**, dashed line).

### Glycan deletion increases mannose trimming throughout the trimer

To elucidate the impact of glycan site mutations at the site-specific level, we performed in-line liquid chromatography-mass spectrometry (LC-MS) analysis of glycopeptides from the above glycan knockout and knock-in constructs. To aid the assignment of glycopeptides, a glycan library was generated by ion-mobility mass spectrometry analysis of an aliquot of unlabeled N-glycans from the BG505 SOSIP.664 protein (**Figure S4, Table 1**). The use of one glycan library for the subsequent analysis of all the glycan site mutants is justified given the likeness of their HILIC-UPLC glycan profiles (**Figure S3**). The LC-MS methodology has been previously validated (Behrens et al., 2016). In addition, we performed glycopeptide analysis on three biological replicates of BG505 SOSIP.664 to confirm that observed differences were not due to experimental variation (**Figure S5**).

The deletion of glycan sites from the network of glycans surrounding the 2G12 epitope generally resulted in increased glycosylation processing at the immediately adjacent sites. This was most significant for the N332T, N411A and N448A glycan deletions, surrounding the N295 glycan site. The loss of each of these glycans resulted in a 61, 80 and 78 percentage point (pp; the arithmetic difference between two percentages) decrease in Man9 at the N295 site, respectively, generally accompanied by a compensatory increase in Man5-8 structures (**Figure 3**). The effect was reciprocal, though less pronounced, with the N295A glycan knockout increasing processing at the N332 and N448 sites (18 and 37 pp decrease in Man9, respectively; **Figure 3**, N295A). Similarly, the N386A mutation resulted in a 42 pp decrease in Man9 at the adjacent N363 site, and the N392A knockout increased processing at the surrounding N339 and N386 sites by 30 and 35 pp, respectively (**Figure 3** N386A).

**Figure 3.**
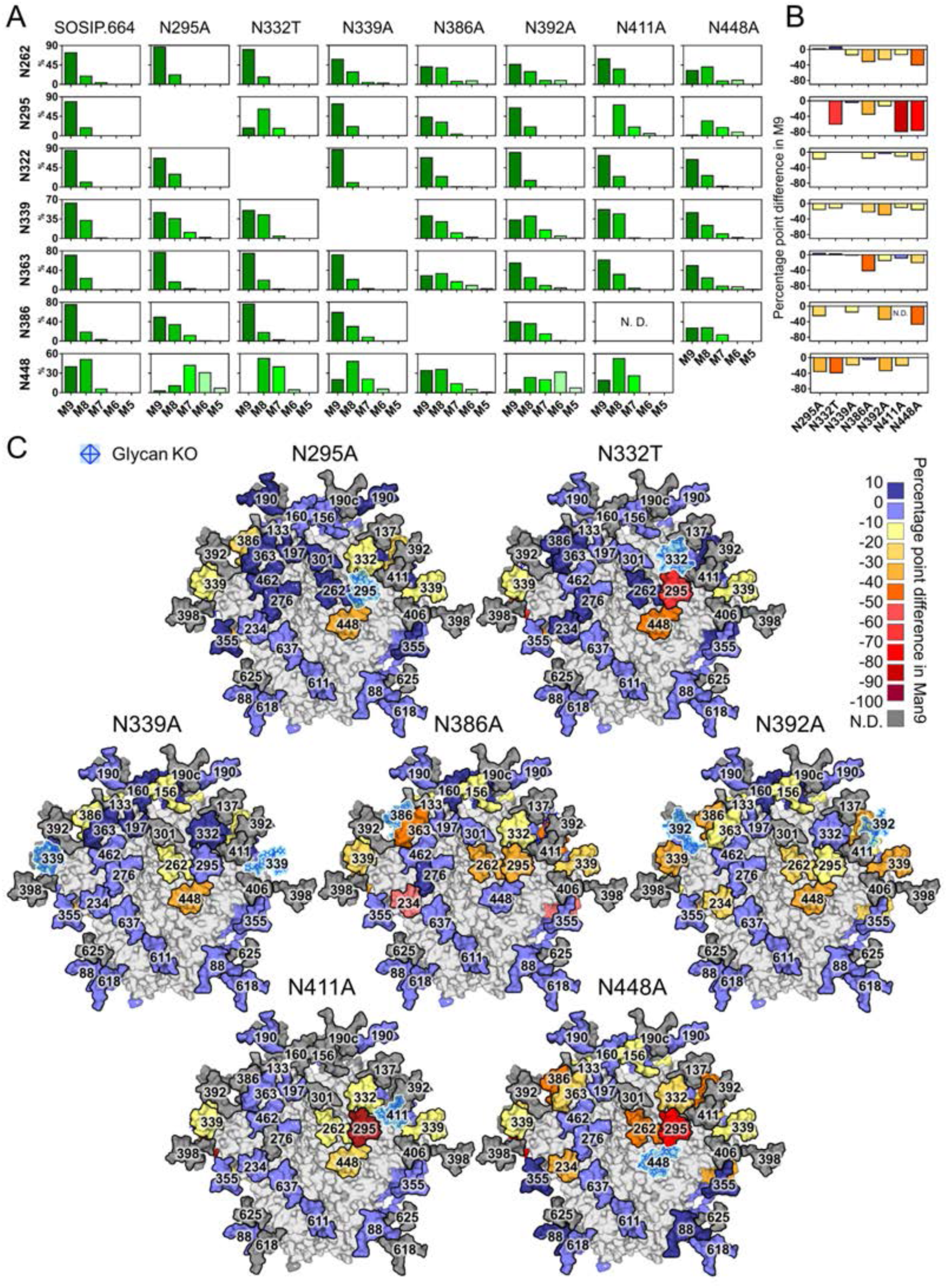
Glycan deletion increases mannose trimming throughout the trimer. A) Relative quantification of IMP sites from BG505 SOSIP.664 and the N295A, N332T, N339A, N386A, N392A, N411A and N448A glycan knockouts. M9 = Man9 (dark green) to M5 = Man5 (pale green). B) The percentage point difference in the abundance of Man9 at IMP sites in the glycan knockouts, compared to BG505 SOSIP.664. Decreases in the abundance of Man9 are colored as per the key in panel C. C) Heat map demonstrating the percentage point difference in the abundance of Man9 at each site in the glycan knockouts, compared to BG505 SOSIP.664. Differences are calculated as follows: (% Man9 in knockout − % Man9 in BG505 SOSIP.664). KO = knockout, N.D. = not determined.

Increased glycosylation processing, however, was not entirely limited to sites adjacent to the glycan knockout. In many instances, glycan site deletion resulted in increased processing emanating from the point of knockout. For example, the N332T knockout also increased processing at the N448 and N339 sites (40 and 13 pp decrease, respectively; **Figure 3**, N332T). As before, the effect appeared reciprocal, as the N448A knockout increased processing at the N332 site (21 pp decrease), located beyond the N295 glycan (**Figure 3**; N448A). Additionally, the N234 glycan, which forms a small cluster with the N276 glycan distal glycan to the IMP, demonstrated increased processing upon the deletion of the N386, N392, or N448 sites (**Figure 3**).

Taken together, the results reveal the varying impact of glycan site knockouts on the processing of glycans across the glycan network, which may relate to underlying structural interactions between the glycans (Gristick et al., 2016; Lemmin et al., 2017; Stewart-Jones et al., 2016).

### Differential effects of glycan additions on processing

In line with high glycan density limiting mannosidase trimming, glycopeptide analysis of the glycan knock-in constructs revealed that the +N241 glycan site addition restricted processing at the neighboring N448 site, resulting in a 21 pp increase in the abundance of Man9 (**Figure 4**).

**Figure 4.**
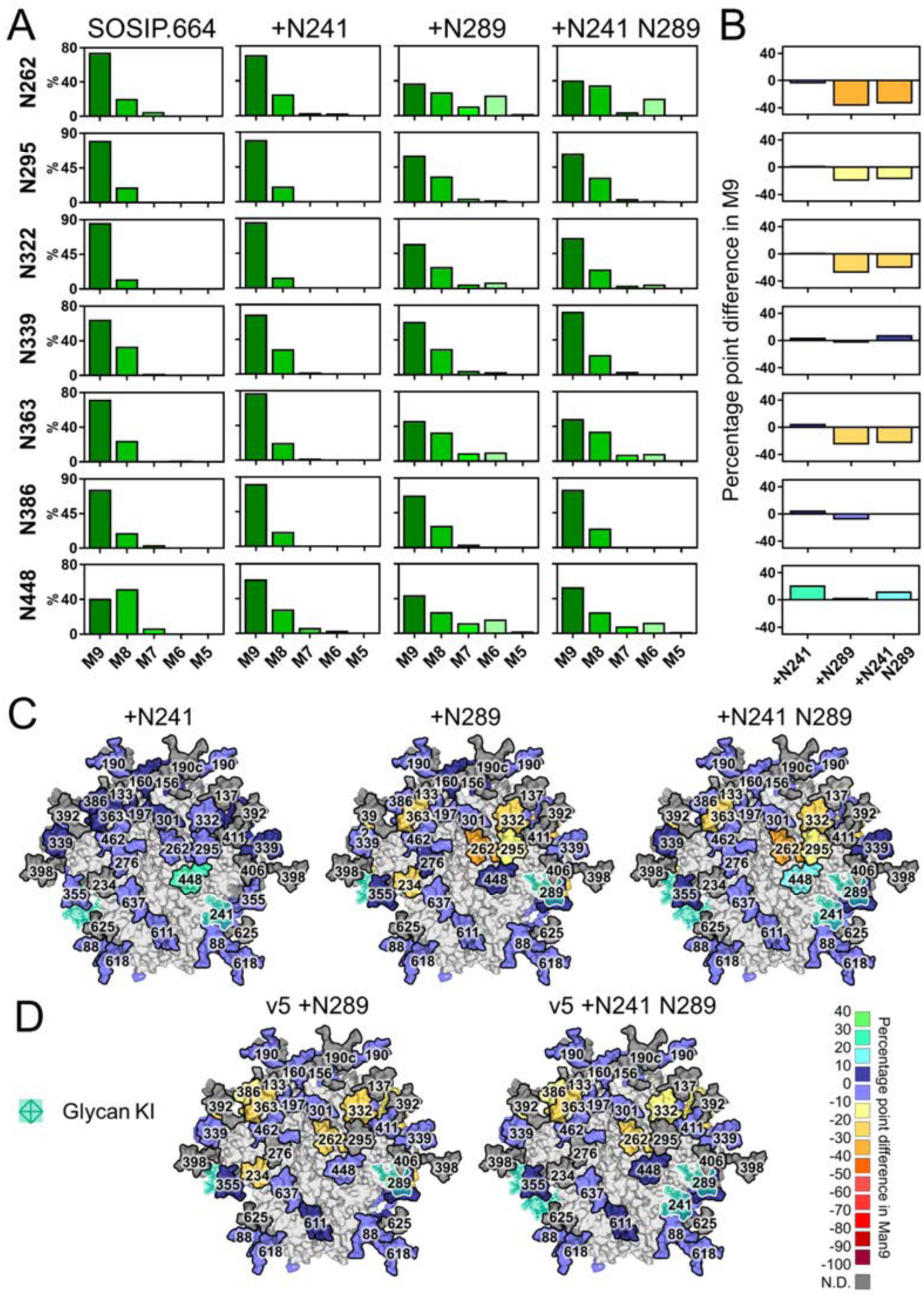
Differential effect of glycan additions on glycosylation processing. A) Relative quantification of IMP sites from BG505 SOSIP.664 and the +N241, +N289 and +N241 N289 glycan knock-ins. M9 = Man9 (dark green) to M5 = Man5 (pale green), complex-type glycans (magenta) are grouped according to their number of antenna (A1-4) and/or presence of a bisecting GlcNAc (B) and/or the presence of core fucose (F) (**Figure S5**). B) The percentage point difference in the abundance of Man9 in the glycan knock-ins, compared to BG505 SOSIP.664. Graphs are colored according to the key in panel D. C) Heat map demonstrating the percentage point difference in the abundance of Man9 at each site in the glycan knock-ins, compared to BG505 SOSIP.664: (% Man9 in knock-in − % Man9 in BG505 SOSIP.664). D) Percentage point difference in the abundance of Man9 at each site in the SOSIP.v5 glycan knock-ins compared to BG505 SOSIP.v5. KI = knock-in.

In contrast, the +N289 site knock-in resulted in increased glycosylation processing, both at sites neighboring the introduced glycan and throughout the trimer. The N262, N295 and N332 sites all displayed a decrease in the abundance of Man9 (37, 20, 28 pp decrease, respectively; **Figure 4**). The N363 and N234 sites were also affected to a similar extent, despite their greater distance from the +N289 knock-in. The +N241 N289 double glycan knock-in displayed both restricted glycosylation processing at the N448 site (12 pp increase in Man9), and increased processing at the N262, N295, N332 and N363 sites (**Figure 4**).

The increased glycan processing associated with the +N289 glycan knock-in is surprising, given that high glycan density is generally associated with restricted glycosylation processing. We had hypothesized that the BG505 SOSIP.664 protein might be unable to accommodate a glycan at this site, and thus the addition of a glycan may be causing wider conformational changes to the protein. We therefore investigated the impact of these glycan knock-ins on the hyperstabilized SOSIP.v5 background (Torrents de la Pena et al., 2017) (**Figure 2A**). In line with the HILIC-UPLC analysis, glycopeptide analysis confirmed that the addition of the N289 glycan to the BG505 SOSIP.v5 background resulted in increased glycan processing, though to a slightly less extent than that observed on the SOSIP.664 background (**Figure 4D**). The SOSIP.v5 +N241 N289 double glycan knock-in also exhibited changes in processing similar to that of the SOSIP.664 background, but not as pronounced.

We had considered that the decrease in the total abundance of oligomannose-type glycans observed by HILIC-UPLC analysis may be, at least partially, explained by the addition of a site(s) comprising predominantly complex-type glycans. However, the precise compositions of the N241 and N289 glycan additions could not be readily determined as they co-occupy peptides with the N234 and N295 sites, respectively. To classify the glycan-type occupying these sites, we subjected the glycopeptides to sequential digests with Endo H (to cleave oligomannose-type glycans) and Peptide-N-glycosidase F (PNGase F; to cleave remaining complex-type glycans) (Cao et al., 2017). Both the N241 and N289 sites were found to be almost exclusively occupied by oligomannose-type glycans, irrespective of SOSIP.664 or SOSIP.v5 background (**Figure S6**), confirming that the observed decreases in the abundance of oligomannose-type glycans, particularly Man9 structures, are solely due to increased processing at other glycan sites on the trimer.

### A network of glycans preserves the PG9 and PG16 epitopes

A large proportion of glycan-targeting bnAbs recognize the glycans of the V1/V2 loops, located at the trimer apex (Walker et al., 2009). This class of bnAbs are typified by the PG9 and PG16 antibodies, which contain very long heavy chain complementarity-determining region 3 (HCDR3), allowing for penetration of the N160 glycan triad at the trimer apex (**Figure 5D**) (McLellan et al., 2011; Pancera et al., 2013). The N160 glycan sits in the center of a network of glycans spanning all three protomers, including the highly conserved N156 and N197 sites (96% and 98%, respectively; **Figure 5B**).

**Figure 5.**
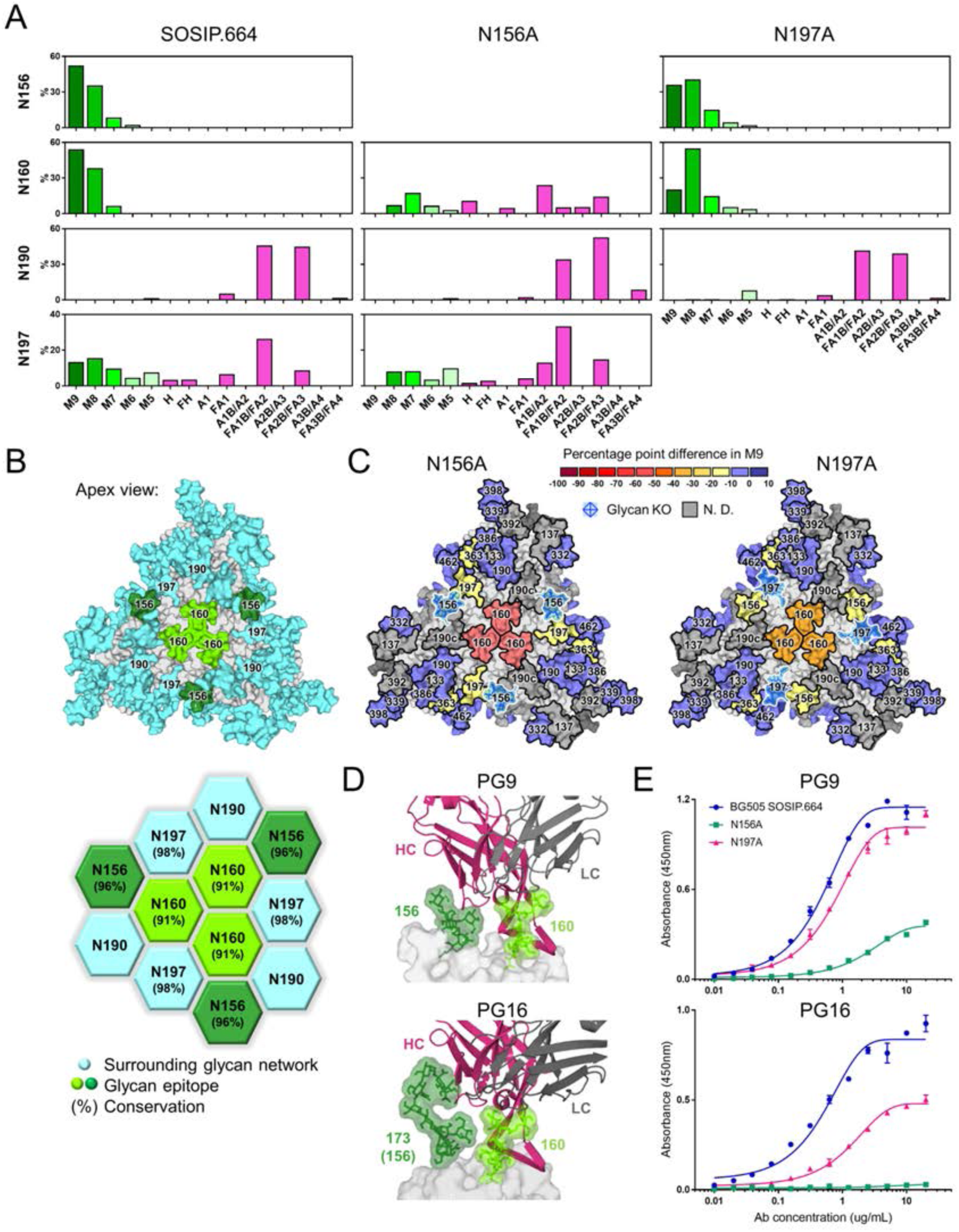
A network of glycans preserves the PG9 and PG16 epitopes. A) Relative quantification of apex glycan sites on BG505 SOSIP.664 and N156A and N197A glycan knockouts. B) Model of a fully glycosylated trimer (top; as described in Figure 1) illustrating the network of glycans at the trimer apex (bottom). (C) Heat maps displaying the percentage point difference in Man9 on N156A and N197A glycan knockouts compared to BG505 SOSIP.664. D) Structure of the PG9 antibody in complex with the V1/V2 region of the CAP45 strain (PDB 3U4E) and PG16 in complex with the V1/V2 region of ZM109 (PDB 4DQO). E) ELISA data of PG9 and PG16 binding to BG505 SOSIP.664 and N156A and N197A glycan knockouts.

In contrast to 2G12, which was largely tolerant of glycan site deletions, the binding of PG16 and, to a lesser extent, PG9 was significantly reduced upon the loss of the N156 glycan (**Figure 5E**). As before, this is somewhat expected as the N156 glycan directly contributes to the antibodies’ epitopes (McLellan et al., 2011; Pancera et al., 2013). However, consistent with previous reports, the N197A glycan knockout also reduced antibody binding (Behrens et al., 2016). Given that the N197 glycan does not contribute to either antibody epitope, we hypothesized that the knockout may be disrupting glycosylation processing at the epitope and performed glycopeptide analysis to address this.

Removal of either the N156 or N197 glycan sites resulted in increased processing at the proximal N160 site (**Figure 5A and C**). This was particularly true of the N156A glycan knockout, which resulted in the complete processing of Man9 to smaller oligomannose-type structures and complex-type glycosylation, such that the dominant peak shifted from Man9 to afucosylated biantennary structures (**Figure 5A**). The N197A glycan knockout resulted in increased oligomannose trimming at the N160 site, resulting in Man8 predominating (**Figure 5A**). The N156A and N197A glycan knockouts also affected each other reciprocally, with each resulting in a slight loss of Man9 at the other site (13.3 and 16.4 pp, respectively; **Figure 5A and B**). We also note that the N363 site exhibited slightly increased oligomannose trimming upon the loss of both glycan sites, though the rest of the trimer appeared unaffected.

## DISCUSSION

There are wide ranging influences of glycan additions and deletions on HIV-1 immune evasion, both in the context of natural infection and in immunization regimens. At one level, the very high density of glycans on Env indicates a selective advantage in using glycans to evade elimination by the host immune system (Scanlan et al., 2007; Wyatt and Sodroski, 1998). However, the glycan shield doesn’t simply evolve to a maximum number of glycans. Instead, the creation and filling of holes during the course of infection illustrates that active rearrangements are required for effective immune evasion (Dacheux et al., 2004; Moore et al., 2012; Wagh et al., 2018; Wei et al., 2003).

In the context of vaccination, the opening and closing of glycan holes in immunogens may prove a useful tool for driving the development of neutralization breadth (Jardine et al., 2013; McGuire et al., 2013; Medina-Ramirez et al., 2017; Ringe et al., 2019; Stamatatos et al., 2017; Steichen et al., 2016). The role of glycans in forming and blocking epitopes is of particular interest as bnAbs can evolve to recognize these structures (Moore et al., 2012). In some instances, the precise processing state of the glycan target is essential for bnAb recognition and neutralization, with glycan heterogeneity manifesting as <100% neutralization plateaus (Doores and Burton, 2010; Kong et al., 2013; McCoy et al., 2015; Pritchard et al., 2015b). The heterogeneity exhibited at a given glycan site can be influenced by the proximity of neighboring glycans (Behrens et al., 2018; Cao et al., 2017; Pritchard et al., 2015a). Using trimeric BG505 SOSIP.664 as a model system, we established the impact of individual glycan site mutations on the glycan networks at two key antigenic regions of the HIV-1 glycan shield, the intrinsic mannose patch and the trimer apex.

The contribution of glycans to many bnAb epitopes has been defined by structural methods, such as X-ray crystallography and cryo-EM, complemented by glycopeptide analysis (Crispin et al., 2018; Ward and Wilson, 2017). In this study, we present the highest resolution structure reported to date of 2G12 in complex with its Env target. We confirm that 2G12 directly contacts the oligomannose-type glycans at four sites and reveal the contributions of the surrounding glycans. It has previously been shown that mutations affecting glycan sites lying outside of bnAb epitopes can disrupt binding and/or neutralization (Behrens et al., 2016; Crispin et al., 2018; McCoy et al., 2016; Scanlan et al., 2002). In line with this, we reported that the deletion of glycan sites in the networks surrounding glycan-dependent antibody epitopes perturbed antibody binding, in some cases to the same extent as mutations directly impacting the epitope. Such observations may be partially explained by disruptions to the fine processing of the glycan epitope upon the mutation of proximal glycan sites, specifically increased trimming by mannosidases.

Despite causing significant disruption to the fine processing of the 2G12 epitope, the N411A knockout displayed increased 2G12 binding, consistent with previous reports (Scanlan et al., 2002). The N411 glycan was unresolved in the cryo-EM structure, though the asparagine residue occupied the middle of 2G12-glycan complex (**Figure S1**). A glycan at this site could potentially result in a clash or the entropy of the glycan could be reduced upon 2G12 binding (**Figure 1**).

The current study was restricted to the analysis of soluble BG505 SOSIP.664 immunogens, an important experimental model for viral glycosylation. We note that soluble SOSIP.664 trimers display somewhat elevated oligomannose levels compared to virion-derived Env (Cao et al., 2018; Struwe et al., 2018). It could, therefore, be argued that mutations that increase the glycan processing of SOSIP.664 trimers are beneficial to generating immunogens that mimic the glycosylation of the native virus. However, oligomannose sites tend to be conserved between SOSIP.664 trimers and virion-derived Env (Cao et al., 2018; Struwe et al., 2018), thus the integrity of the mannose patch is associated with well folded trimers. For example, within the BG505 SOSIP.664 experimental system, substantial changes in the abundance of oligomannose-type glycans can indicate deviations away from native-like conformations (Pritchard et al., 2015c).

In this context, we note that the mutation of N-linked glycan sites can induce glycoprotein misfolding or conformational changes (Kong et al., 2015; Pritchard et al., 2015a; Sanders et al., 2002; Wang et al., 2013). Here, the glycan mutants were purified using a quaternary structure-dependent antibody affinity step, PGT145 or PGT151 (Blattner et al., 2014; Walker et al., 2011), to minimize the contribution of misfolded proteins to the analysis. Accordingly, all the mutants displayed a glycosylation profile dominated by oligomannose-type glycans, a signature of native-like trimer configuration (Behrens and Crispin, 2017; Behrens et al., 2017a; Pritchard et al., 2015c).(Cao et al., 2017; Pritchard et al., 2015c)However, the unexpected increase in glycosylation processing observed upon the knock-in of the N289 glycan, in both the SOSIP.664 trimers and hyperstabilized SOSIP.v5 trimers, suggests that this mutation may be causing a degree of protein instability. We note that the +N289 knock-in was generated by mutating a proline at the 291 position. The results highlight the importance of characterizing the glycosylation of all candidate immunogens in depth (Behrens et al., 2017b).

Understanding the interdependence of glycans and their processing states is important in revealing how viral mutations can influence distant epitopes. Similarly, in immunogen design, the presence or absence of holes in the glycan shield could have wider antigenic and immunogenic consequences through glycan-glycan network effects. While the processing state of some glycans is key to the formation of bnAb epitopes (Kong et al., 2013; Pritchard et al., 2015b), the network may be dominated simply by the presence or absence of glycans independent of their processing state. Consistent with this view, we report that both the deletion of glycan sites surrounding the 2G12 epitope, and the addition of the N289 glycan, increased the mannosidase trimming of the epitope. However, only glycan site knockouts had a measurable impact on 2G12 binding. Thus, the epitope is likely maintained by the network of glycans surrounding the epitope providing structural support to the four glycans directly involved in 2G12 binding(Dunlop et al., 2010).

The same network effects appear to apply to the epitopes of the apex binding antibodies, PG9 and PG16. Previous studies have reported the dependence of apex targeting bnAbs on sialylated glycans at the N156 site (Andrabi et al., 2017; McLellan et al., 2011; Pancera et al., 2013). However, glycopeptide analyses presented both here and previously, report this site to be predominantly occupied by oligomannose-type glycans (Behrens et al., 2017a; Behrens et al., 2016; Cao et al., 2017; Cao et al., 2018). Thus, these bnAbs are able to tolerate glycan heterogeneity. The considerable loss of binding upon glycan site deletion is therefore symptomatic of the structural role the network of surrounding glycans plays in stabilizing this glycan epitope. At the apex in particular, the impact of the deletion of an individual glycan site may be amplified three-fold.

It is not yet known the extent to which a successful vaccine candidate must display precise glycan epitopes. However, the results presented here shed light on the role of individual glycan sites in the fine processing of bnAb epitopes. We reveal the hierarchical role of glycan networks in stabilizing the structure and fine processing of two key bnAb epitopes, the trimer apex and intrinsic mannose patch. A growing understanding of the factors shaping the glycosylation of Env will aid the continued development of HIV-1 immunogens.

## Author contributions

G.E.S, C.A.C, M.J.v.G, A.D, A-J.B, A.M, N.D.V and S.V performed experimental work. G.E.S, C.A.C, Y.W and J.D.A analyzed the data. G.E.S and M.C wrote the paper. R.W.S, A.B.W and M.C designed the study. All authors read and approved the final manuscript.

## Acknowledgments

This project has received funding from the European Union’s Horizon 2020 Research and Innovation programme under grant agreement No. 681137 (to R.W.S. and M.C), the Bill and Melinda Gates Foundation through the Collaboration for AIDS Vaccine Discovery (OPP1084519 and OPP1115782; to M.C., and OPP1132237; to R.W.S.), the U.S. National Institutes of Health Grant P01 AI110657 (to A.B.W. and R.W.S.), the Center for HIV/AIDS Vaccine Immunology and Immunogen Discovery (1UM1AI100663; to M.C.). C.A.C is supported by NIH F31 Ruth L. Kirschstein Predoctoral Award Al131873 and by the Achievement Rewards for College Scientists Foundation. M.J.v.G. is a recipient of a Mathilde Krim Fellowship from the American Foundation for AIDS Research, (amfAR; grant 109514-61-RKVA). R.W.S. is a recipient of a Vici grant from the Netherlands Organization for Scientific Research (NWO).

## Declaration of Interests

The International AIDS Vaccine Initiative (IAVI) has previously filed a patent relating to the BG505 SOSIP.664 trimer: U.S. Prov. Appln. No. 61/772,739, entitled ‘‘HIV-1 Envelope Glycoprotein,’’ with R.W.S. and A.B.W. amongst the co-inventors, but no patents have been filed on any work described here.

## STAR methods

### CONTACT FOR REAGENT AND RESOURCE SHARING

Further information and requests for resources and reagents should be directed to and will be fulfilled by the Lead Contact, Max Crispin

### EXPERIMENTAL MODEL AND SUBJECT DETAILS

#### HEK 293F cell culture

HEK 293F cells were maintained at a density of 0.1-3×10^6^ cells per mL at 37°C and 125 rpm shaking. The cells were transfected at a density of 1×10^6^ cells per mL and incubated for 5 days at 37°C with 8% CO_2_ and 125 rpm shaking.

### METHOD DETAILS

#### BG505 SOSIP.664 and SOSIP.v5 construct design

Asparagine residues within the N295, N339, N386, N392, N411 and N448 consensus sequence, N-X-S/T (where X ≠ P), were mutated to alanine. For the N332 site, we reverted the asparagine to threonine, as per the parental BG505 sequence. To introduce N-linked glycan sites at the N241 and N289 sites, the serine at the 241 position was mutated to asparagine, and/or the proline at the 291 position was mutated to serine. Mutants were created using the QuikChange II site-directed mutagenesis kit (Agilent), as per the manufacturer’s instructions.

#### Expression and purification of proteins

BG505 SOSIP.664 and SOSIP.v5 proteins were transiently (co-)expressed in HEK 293F cells with a Furin expression plasmid at a ratio of (4:1). BG505 SOSIP.664 and SOSIP.v5 proteins were purified using either 2G12, PGT145, or PGT151 affinity chromatography, as previously described (Sanders et al., 2013). Comparisons were only drawn between BG505 SOSIP.664 proteins purified using the same antibody.

PGV04 Fab and 2G12, PGT145, PGT151, PG16 and PG9 IgG were produced by co-expression of the heavy and light chain genes in HEK293F cells. PGV04 Fab was purified using KappaSelect and Mono-S columns (GE Healthcare). 2G12, PGT145, PGT151, PG16 and PG9 IgG were purified with a Protein A column (GE Healthcare). 2G12 Fab_2_ was prepared by digesting 2G12 IgG with papain using a previously described protocol (Calarese et al., 2003).

#### Cryo-electron microscopy and model building

BG505 SOSIP.664 purified by 2G12 affinity chromatography was mixed with 10-fold molar excess PGV04 Fab and 2G12 Fab_2_ and incubated overnight at room temperature. The trimer/Fab complex was purified by size exclusion chromatography using a Superose 6 column (GE Healthcare). The fractions containing the complex were pooled and concentrated using a 100-kDa concentrator (Amicon Ultra, Millipore) to ∼40 μL at 2.5 mg/mL. 5 μL of the complex was incubated with 3 μL of a fresh n-Dodecyl-B-D-Maltoside solution at 1.8 mM. A 3 μL aliquot of the complex/detergent mix was applied to a C-Flat grid (CF-2/2-4C, Electron Microscopy Sciences, Protochips, Inc.) which had been plasma cleaned for 5 seconds using a mixture of Ar/O_2_ (Gatan Solarus 950 Plasma system). The sampled was manually blotted off, and then immediately plunged into liquid ethane using a manual freeze plunger.

Movies were collected via the Leginon interface on a FEI Titan Krios operating at 300 keV mounted with a Gatan K2 direct electron detector (Carragher et al., 2000). Each movie was collected in counting mode at 22,500 × nominal magnification resulting in a calibrated pixel size of 1.31 Å/pix at the object level. A dose rate of ∼10 e^-^/(pix*s) was used; exposure time was 200 ms per frame. The data collection resulted in a total of 2,184 movies containing 50 frames each. Total dose per movie was 76 e^-^/Å^2^. Data were collected with a defocus range of −1.5 to −3.0 microns. Movies were imported into cryoSPARC v2 and frames were aligned using full-frame motion correction (Punjani et al., 2017). The contrast transfer function (CTF) for each aligned micrograph was estimated using Gctf (Zhang, 2016). The HIV Env portion of PDB 5ACO was converted to an EM density and low pass filtered to 40 Å using pdb2mrc and subsequently used as a template for particle picking within cryoSPARC v2 (Lee et al., 2015; Ludtke et al., 1999; Punjani et al., 2017). 2D classification, Ab-initio 3D reconstruction, homogenous 3D refinement, and local motion correction were conducted with cryoSPARC v2 (Punjani et al., 2017). Per-particle CTF estimation was conducted using using Gctf (Zhang, 2016). The final C3 symmetric, 3.8 Å reconstruction was obtained using non-uniform refinement within cryoSPARC v2 (Punjani et al., 2017). The per gp120 protomer occupancy of the PGV04 Fab was low and resulted in weak density when the EM data were refined using C3 symmetry. As such, the PGV04 Fab was not included during the model building process.

An initial model was made by docking the gp120 and gp41 domains from the BG505 SOSIP.664 structure (PDB 5ACO) and the 2G12 Fab_2_ structure (PDB 6N2X) into the EM density map using UCSF Chimera (Calarese et al., 2003; Lee et al., 2015; Pettersen et al., 2004). The resulting model was symmetrically refined into the EM density map using RosettaRelax (DiMaio et al., 2015). Glycans were built manually using the Carbohydrate module in Coot and refined into the EM density map using Rosetta (Emsley and Crispin, 2018; Frenz et al., 2019). Model accuracy and fit-to-map were assessed using Molprobity, EMRinger, Privateer, CARP, and pdb-care (Agirre et al., 2015; Barad et al., 2015; Lutteke et al., 2005; Williams et al., 2018). Binding surface area calculations were performed using jsPISA (Krissinel, 2015).

#### Enzymatic release of N-linked glycans

N-linked glycans were released from BG505 SOSIP.664 proteins by in-gel digestion with PNGase F (New England Biolabs). Proteins were resolved by SDS–PAGE and stained with Coomassie Blue. Following destaining, the protein band was excised and washed alternatively with acetonitrile and water. Gel bands were then incubated with PNGase F for 16□h at 37□°C. Released glycans were eluted from the gel with water and dried in a SpeedVac concentrator.

#### Fluorescent labeling of N-linked glycans

Released glycans were fluorescently labeled with procainamide (Abcam). Dried glycans were resuspended in 30□μL water before addition of 80□μL labeling mixture: 110 mg/mL procainamide, 60 mg/mL sodium cyanoborohydride in a solution of 70% dimethyl sulfoxide, 30% acetic acid. Samples were incubated at 65□°C for 4 h. Labeled glycans were purified using Spe-ed Amide-2 cartridges (Applied Separations).

#### HILIC-UPLC analysis of the N-linked glycans

Fluorescently labeled glycans were separated by HILIC-UPLC on a Waters ACQUITY H-Class instrument using a 2.1□mm × 10□mm Glycan BEH Amide Column (1.7□μm particle size; Waters). The following gradient was run: time=0□min (t=0): 22% A, 78% B (flow rate of 0.5□mL/min); t=38.5: 44.1% A, 55.9% B; t=39.5: 100% A, 0% B (0.25□mL/min); t=44.5: 100% A, 0% B; t=46.5: 22% A, 78% B (0.5 mL/min), where solvent A was 50□mM ammonium formate, pH 4.4, and solvent B was acetonitrile. Fluorescence was measured using an excitation wavelength of 310□nm and a detection wavelength of 370□nm.

#### Oligomannose-type glycan quantification

Quantification of oligomannose-type glycans was measured by digestion with Endo H, which cleaves oligomannose-(and hybrid-) type glycans, but not complex-type (New England Biolabs). Labeled glycans were resuspended in water and digested with Endo H for 16□h at 37□°C. Digested glycans were cleaned using a PVDF protein-binding membrane plate (Merck Millipore) prior to HILIC-UPLC analysis as above. The abundance of oligomannose-type glycans was calculated, as a relative percentage, by integration of the HILIC-UPLC chromatograms before and after Endo H digestion, following normalization.

#### Ion mobility-mass spectrometry of N-linked glycans

To guide subsequent glycopeptide analyses, we performed IM-MS on a separate, unlabeled aliquot of PNGase F-released glycans from the BG505 SOSIP.664 protein. Glycan compositions were determined using traveling wave IM-MS measurements performed on a Synapt G2Si instrument (Waters, Manchester, UK). The glycan sample was cleaned with a Nafion 117 membrane and a trace amount of ammonium phosphate was added to promote phosphate adduct formation. Glycans were analyzed by nano-electrospray with direct infusion using a Synapt G2Si instrument (Waters, Manchester, UK) with the following settings: capillary voltage, 0.8–1.0 kV; sample cone, 100 V; extraction cone, 25 V; cone gas, 40 L/h; source temperature, 150 °C; trap collision voltage, 4–160 V; transfer collision voltage, 4 V; trap direct current bias, 35–65 V; IMS wave velocity, 450 m/s; IMS wave height, 40 V; trap gas flow, 2 mL/min; IMS gas flow, 80 mL/min. Data were acquired and processed with MassLynx v4.1 and Driftscope version 2.8 software (Waters, Manchester, UK). Structural assignments were based on previously described IM-MS of BG505 SOSIP.664 glycans (Behrens et al., 2016).

#### Reduction, alkylation and digestion of Env proteins

BG505 SOSIP.664 and SOSIP.v5 proteins (100-150 ug each) were denatured, reduced, and alkylated by sequential 1 h incubations at room temperature (RT) in the following solutions: 50 mM Tris/HCl, pH 8.0 buffer containing 6 M urea and 5 mM dithiothreitol (DTT), followed by the addition of 20 mM iodacetamide (IAA) for a further 1h at RT in the dark, and then additional DTT (20 mM), to eliminate residual IAA. The proteins were then buffer-exchanged into 50 mM Tris/HCl, pH 8.0 using Vivaspin columns and aliquots were digested with trypsin or chymotrypsin (Mass Spectrometry Grade, Promega) at a ratio of 1:30 (w/w) for 16 h at 37 °C. The reactions were dried and glycopeptides were extracted using C18 Zip-tip (Merck Millipore) following the manufacturer’s protocol.

#### Liquid chromatography-mass spectrometry analysis of glycopeptides

Eluted glycopeptides were dried again and re-suspended in 0.1% formic acid prior to mass spectrometry analysis. An aliquot of glycopeptides was analyzed by LC-MS with an Easy-nLC 1200 system coupled to an Orbitrap Fusion mass spectrometer (Thermo Fisher Scientific) using higher energy collisional dissociation (HCD) fragmentation. Peptides were separated using an EasySpray PepMap RSLC C18 column (75 μm x 75 cm) with a 275 minute linear gradient consisting of 0%–32% acetonitrile in 0.1% formic acid over 240 minutes followed by 35 minutes of 80% acetonitrile in 0.1% formic acid. The flow rate was set to 200 nL/min. The spray voltage was set to 2.8 kV and the temperature of the heated capillary was set to 275 °C. HCD collision energy was set to 50%, appropriate for fragmentation of glycopeptide ions. Glycopeptide fragmentation data were extracted from the raw file using Byonic™ (Version 2.7) and Byologic™ software (Version 2.3; Protein Metrics Inc.). The glycopeptide fragmentation data were evaluated manually for each glycopeptide; the peptide was scored as true-positive when the correct b and y fragment ions were observed along with oxonium ions corresponding to the glycan identified. The relative abundance of each glycoform at each site was calculated using the extracted ion chromatograms for true-positive peptides.

#### Site-specific glycan classification

Remaining glycopeptides were first digested with Endo H to cleave oligomannose- and hybrid-type glycans, leaving a single GlcNAc residue at the corresponding site. The reaction mixture was then dried completely and resuspended in a mixture containing 50 mM ammonium bicarbonate and PNGase F using only O^18^-labeled water (Sigma-Aldrich) throughout. This second reaction cleaves the remaining complex-type glycans, leaving the GlcNAc residues intact. The use of H_2_O^18^ in this reaction enables complex glycan sites to be differentiated from unoccupied glycan sites as the hydrolysis of the glycosidic bond by PNGase F leaves an O^18^ isotope on the resulting aspartic acid residue. The resultant peptides were purified by C18 ZipTip, as outlined above, and subjected to LC-MS in a similar manner to before, but using a lower HCD energy of 27% as glycan fragmentation was not required. Data analysis was performed as above.

#### ELISAs

Antibody 2G12 was expressed in HEK 293F cells (1:2 ratio of heavy to light chain plasmids) and purified by Protein A affinity chromatography according to the manufacturer’s instructions (GE Healthcare). High binding 96 well assay plates (Corning) were incubated with BG505 SOSIP.664 proteins (10 ug/mL in PBS) overnight at 4 °C. Plates were washed with a solution of PBS containing 0.5% Tween 20 (v/v) and blocked for 1h at RT with 5% milk in PBS + 0.5% Tween. After another wash step, the primary antibody was incubated (1:2 dilution series with a starting concentration of 20 ug/mL) in PBS for 1h at RT. Plates were washed and an anti-human IgG conjugated to Horseradish Peroxidase (Abcam) secondary antibody was added at a 1:2’000 dilution in PBS. Plates were washed and TMB substrate solution (Thermo Fisher Scientific) was added. The reaction was stopped with sulfuric acid after 5□min and the OD 450□nm was measured.

### QUANTIFICATION AND STATISTICAL ANALYSIS

The integration of peaks corresponding to fluorescently labeled N-glycans was performed using Empower 3.0 (Waters, Manchester, UK) (Figure 2 and S3). The IM-MS data used to generate the glycan library were acquired and processed with MassLynx v4.1 and Driftscope version 2.8 software (Waters, Manchester, UK) (Figure S4). Chromatographic areas were extracted for site-specific analysis using Byonic™ (Version 2.7) and Byologic™ software (Version 2.3) by Protein Metrics (Figure 3, 4, 5, S5 and S6).

### DATA AND SOFTWARE AVAILABILITY

The cryo EM map and atomic model of BG505 SOSIP.664 with 2G12 Fab_2_ was deposited with the Electron Microscopy Data Bank and the Protein Data Bank under accession codes.

