## Supplemental Data for "Networks of HIV-1 envelope glycans maintain antibody epitopes in the face of glycan additions and deletions"

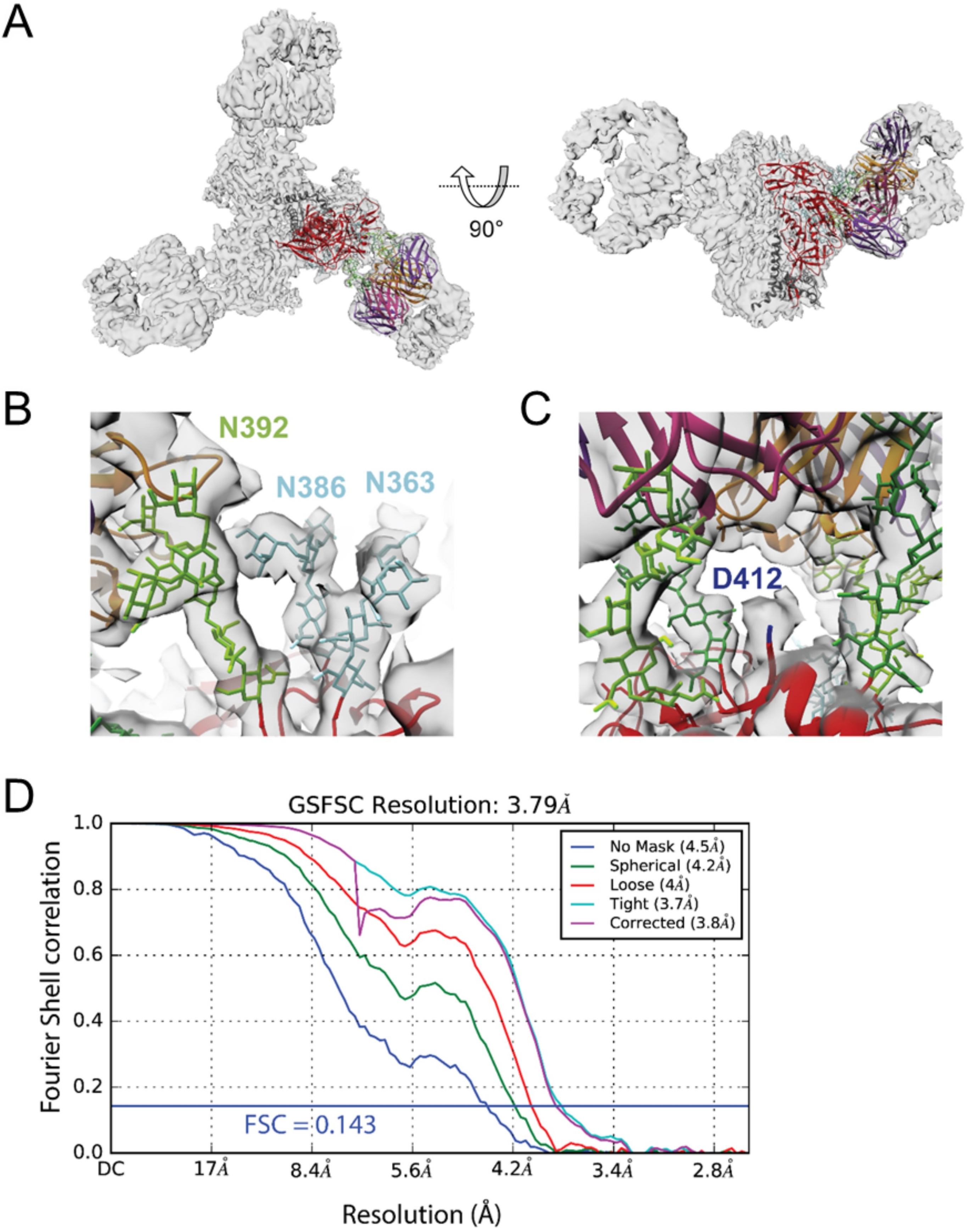


**Figure S1. Cryo-EM of BG505 SOSIP.664 and 2G12 Fab$_2$.** Related to Figure 1. A) 3.8 Å cryo-EM 3D reconstruction and refined atomic model of BG505 SOSIP.664 with 2G12 Fab2. B) The close proximity of the N363 and N386 glycans to the N392 glycan could provide support for 2G12 binding via glycan/glycan interactions. C) No coordinated density was seen for the N411 amino acid or the associated N-linked glycan. D) Fourier shell correlation (FSC) curves calculated in cryoSPARC during final non-uniform refinement.

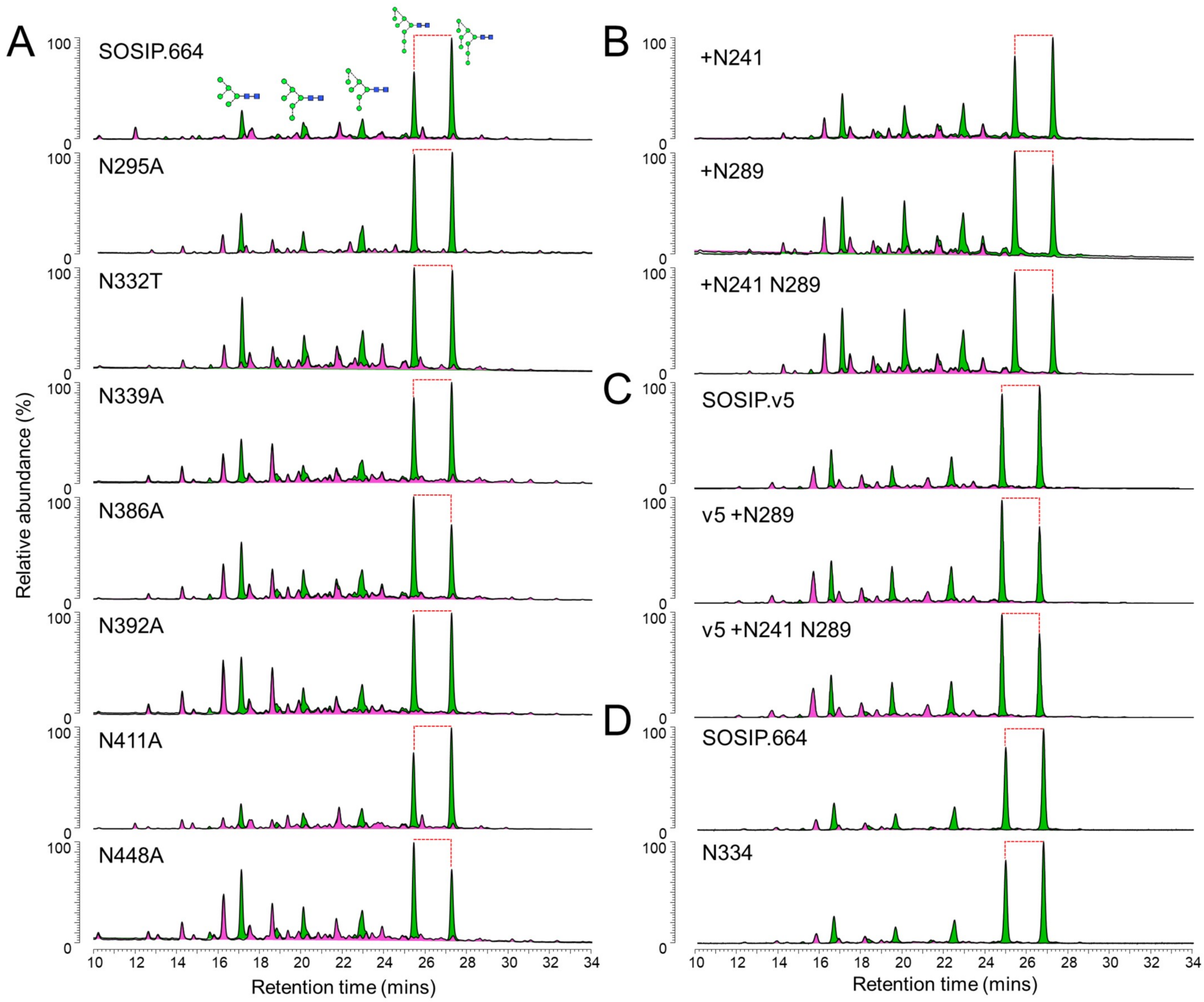


**Figure S2. Glycosylation profiles of glycan mutants.** Related to Figure 2. Hydrophilic interaction liquid chromatography-ultra performance liquid chromatography of PNGase F-released N-linked glycans from BG505 SOSIP.664 and SOSIP.v5 glycan mutants. Oligomannose-type glycans (judged by sensitivity to Endo H digestion) are highlighted in green. Complex-type glycans are shown in pink.

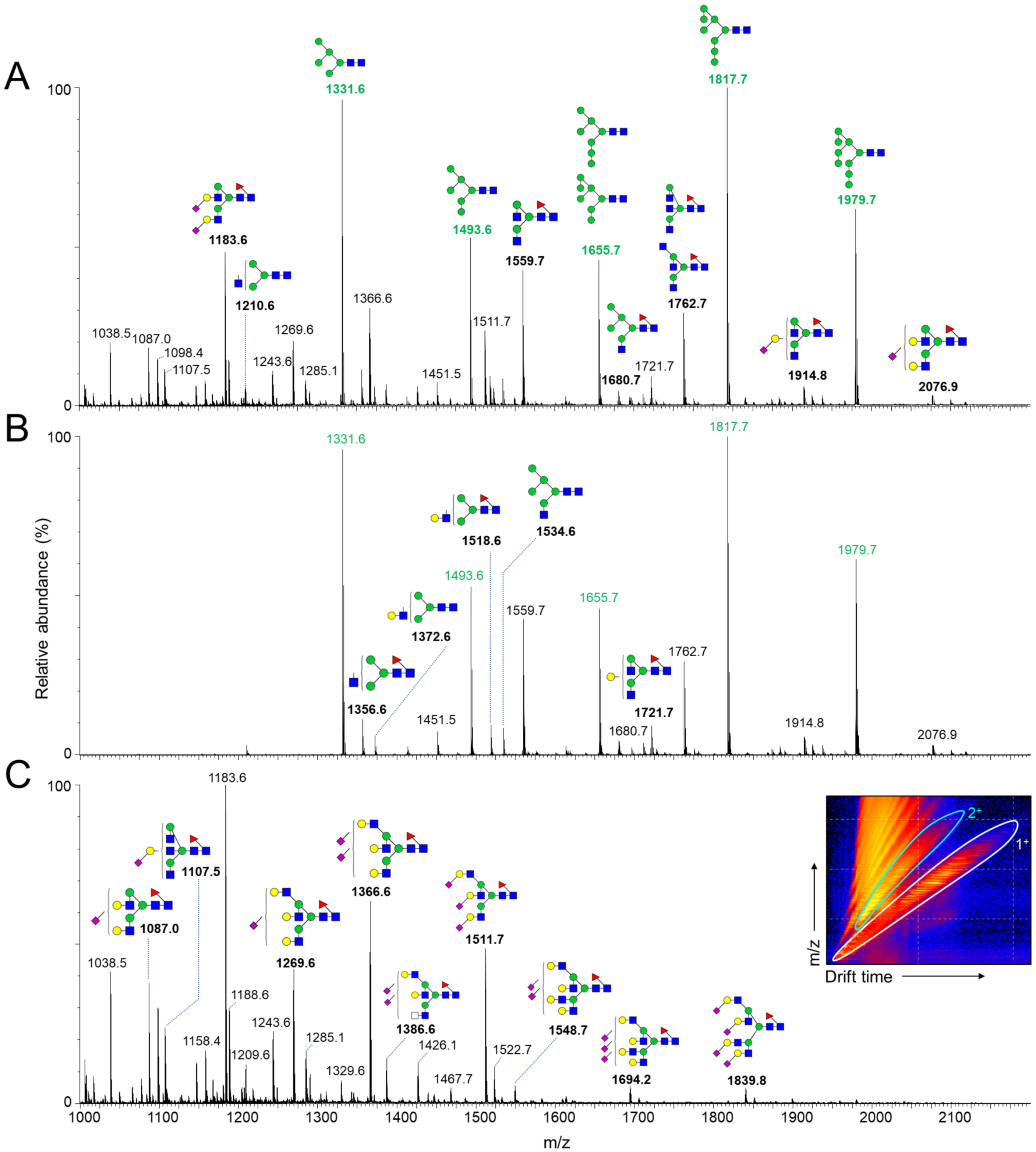


**Figure S3. Generation of a glycan library.** Related to Figures 3, 4 and 5. A) A glycan library was generated by negative ion electrospray ion mobility-mass spectrometry of an aliquot of unlabelled N-glycans from BG505 SOSIP.664 trimers. B) Mobility-extracted singly charged negative ions. C) Mobility-extracted doubly charged negative ions. The corresponding ions are encircled in white and blue, respectively, in the inset ion mobility drift plot.

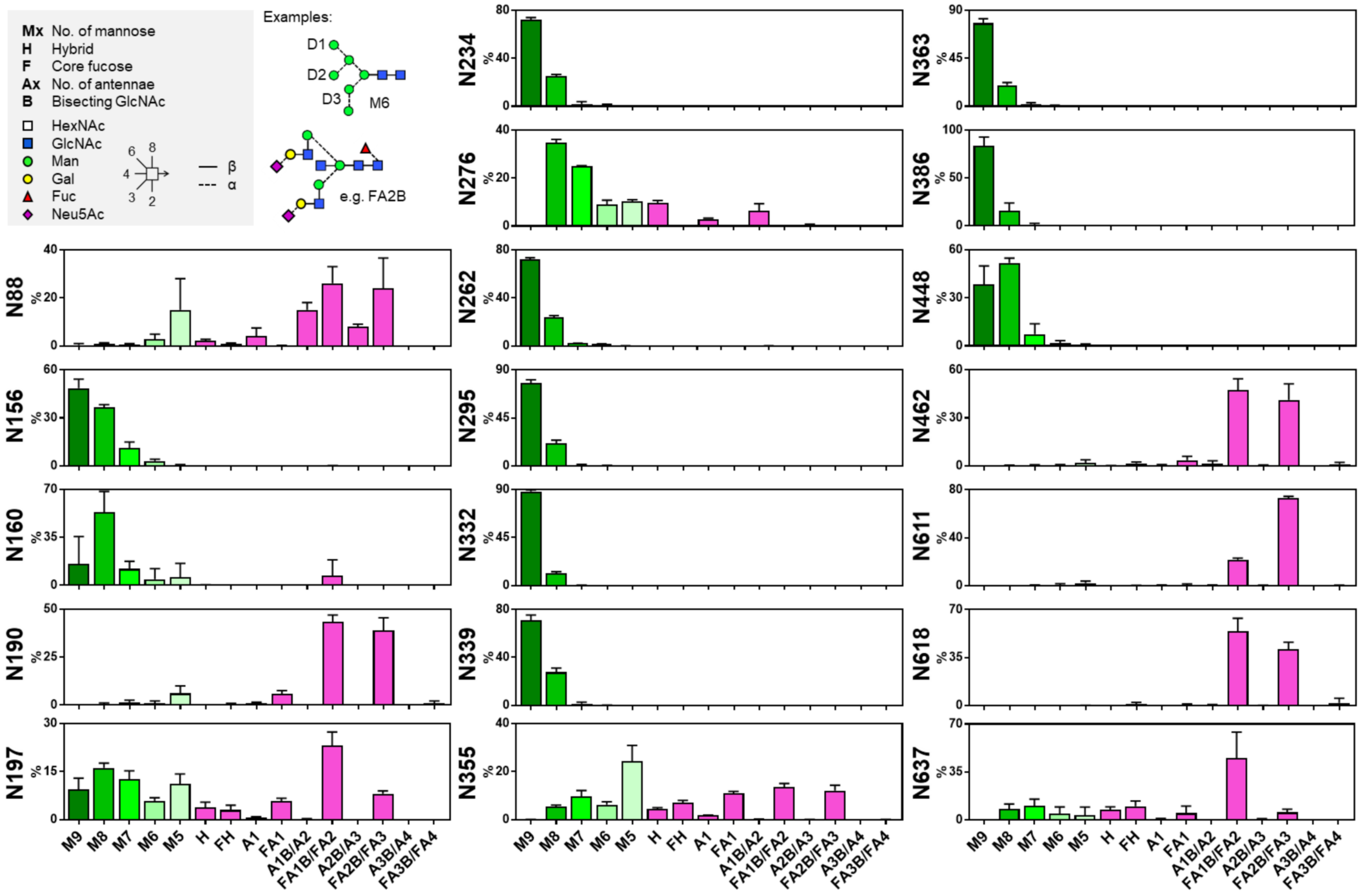


**Figure S4. Reproducibility of LC-MS glycopeptide analysis.** Related to Figures 3, 4 and 5. Relative quantification of glycan sites from three biological replicates of BG505 SOSIP.664. Error bars represent the standard error of the mean. The inset key depicts the glycan nomenclature and classification system used throughout this manuscript. HexNAc = N-acetylhexosamine, GlcNAc = N-acetylglucosamine, Man = mannose, Gal = galactose, Fuc = fucose, Neu5Ac = sialic acid. Glycan nomenclature according to the Consortium for Functional Glycomics, with linkages as per the Oxford system.

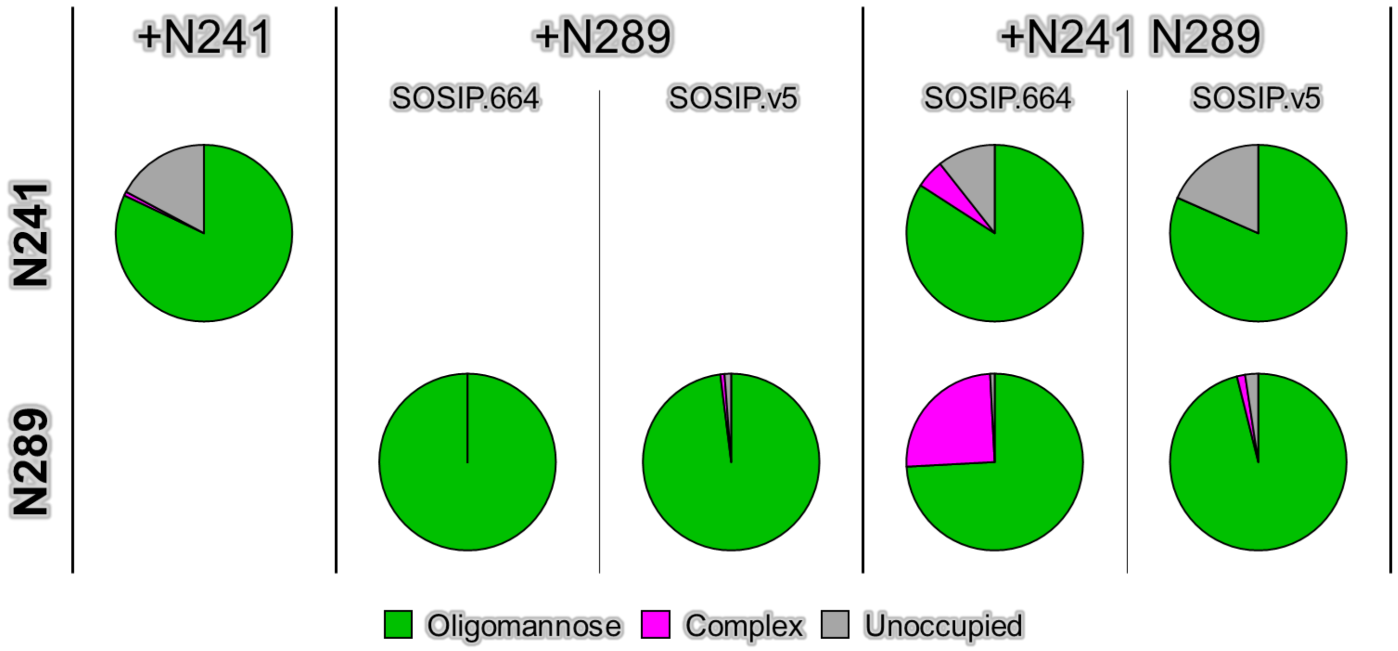


**Figure S5. Classification of glycan-type at the N241 and N289 sites.** Related to Figure 4. Relative quantification of the glycan-type occupying the N241 and N289 sites in the SOSIP.664 and SOSIP.v5 knock-in mutants. Glycopeptides were digested with Endo H to cleave oligomannose-type glycans, leaving behind a GlcNAc residue. The remaining complex-type glycans were then digested with PNGase F in the presence of $O^{18}$-labeled water, to convert the Asn to an $O^{18}$-labeled Asp, prior to LC-MS analysis.

**A**

| | N88 | N130 | N133 | N136 | N137 | N141 | N156 | N160 | N185e | N187 | N188 | N190 | N190c | N197 | N230 | N234 | N241 | N262 | N276 | N289 | N295 | N301 | N332 | N339 | N355 |
|---|---|---|---|---|---|---|---|---|---|---|---|---|---|---|---|---|---|---|---|---|---|---|---|---|---|
| A | 99 | 47 | 14 | 11 | 22 | 17 | 98 | 94 | 7 | 33 | 6 | 4 | 12 | 99 | 3 | 80 | 96 | 100 | 97 | 22 | 54 | 94 | 66 | 63 | 5 |
| B | 99 | 73 | 10 | 23 | 25 | 17 | 96 | 89 | 6 | 22 | 45 | 2 | 8 | 98 | 26 | 72 | 97 | 100 | 97 | 67 | 79 | 95 | 85 | 70 | 4 |
| C | 98 | 54 | 28 | 9 | 25 | 12 | 95 | 92 | 6 | 41 | 8 | 2 | 8 | 98 | 78 | 84 | 97 | 98 | 92 | 79 | 20 | 95 | 77 | 71 | 4 |
| D | 98 | 71 | 3 | 22 | 32 | 9 | 91 | 94 | 10 | 34 | 38 | 3 | 17 | 98 | 17 | 84 | 95 | 99 | 91 | 87 | 76 | 73 | 80 | 75 | 1 |
| F | 98 | 52 | 13 | 10 | 25 | 16 | 95 | 90 | 6 | 29 | 8 | 2 | 11 | 98 | 10 | 83 | 95 | 100 | 98 | 94 | 95 | 95 | 75 | 79 | 6 |
| G | 98 | 62 | 27 | 11 | 28 | 11 | 99 | 94 | 4 | 27 | 23 | 2 | 21 | 99 | 4 | 80 | 97 | 99 | 97 | 79 | 48 | 98 | 70 | 16 | 5 |
| AE | 98 | 66 | 14 | 3 | 14 | 16 | 98 | 93 | 8 | 39 | 6 | 2 | 6 | 98 | 0 | 86 | 99 | 100 | 95 | 96 | 85 | 79 | 2 | 47 | 4 |
| Total | 99 | 64 | 16 | 15 | 24 | 15 | 96 | 91 | 6 | 31 | 26 | 2 | 9 | 98 | 37 | 78 | 97 | 99 | 95 | 72 | 60 | 93 | 72 | 67 | 4 |

| | N356 | N362 | N363 | N386 | N392 | N396 | N398 | N406 | N407 | N411 | N442 | N448 | N461 | N462 | N611 | N616 | N618 | N625 | N637 |
|---|---|---|---|---|---|---|---|---|---|---|---|---|---|---|---|---|---|---|---|
| A | 82 | 22 | 42 | 85 | 82 | 22 | 39 | 5 | 5 | 51 | 6 | 93 | 27 | 30 | 98 | 95 | 2 | 95 | 96 |
| B | 82 | 59 | 5 | 83 | 83 | 32 | 38 | 7 | 6 | 42 | 4 | 92 | 22 | 30 | 99 | 89 | 7 | 97 | 97 |
| C | 74 | 19 | 3 | 88 | 72 | 24 | 27 | 4 | 4 | 44 | 73 | 74 | 18 | 29 | 98 | 92 | 4 | 97 | 97 |
| D | 17 | 10 | 3 | 91 | 79 | 22 | 20 | 7 | 6 | 38 | 2 | 98 | 27 | 39 | 99 | 95 | 2 | 94 | 93 |
| F | 84 | 62 | 0 | 84 | 73 | 2 | 8 | 0 | 0 | 65 | 43 | 71 | 16 | 37 | 100 | 95 | 0 | 100 | 95 |
| G | 32 | 52 | 0 | 92 | 68 | 8 | 22 | 0 | 0 | 49 | 59 | 87 | 28 | 32 | 100 | 95 | 2 | 97 | 98 |
| AE | 75 | 1 | 0 | 93 | 77 | 15 | 43 | 1 | 1 | 64 | 7 | 95 | 31 | 25 | 98 | 98 | 0 | 97 | 98 |
| Total | 76 | 37 | 6 | 86 | 78 | 26 | 34 | 5 | 5 | 46 | 27 | 87 | 22 | 29 | 99 | 92 | 5 | 97 | 97 |

**B**

| | Mean |
|---|---|
| A | 28.9 |
| B | 30.5 |
| C | 29.1 |
| D | 29.2 |
| F | 29.7 |
| G | 30.0 |
| AE | 28.6 |

0 — 100 %

**Figure S6. Conservation of PNGS across clades.** Related to Figures 1 and 5. A) Percentage conservation of individual PNGS. B) Mean number of total PNGS. The analysis was performed using sequence data obtained from the 2017 filtered web alignment from the Los Alamos HIV sequence database (https://www.hiv.lanl.gov). Clade A, *n* = 338; clade B, *n* = 2351; clade C, *n* = 1524; clade D, *n* = 127; clade F, *n* = 63; clade G, *n* = 114, clade AE, *n* = 535. Only the second position Asn was included for NN[S/T][S/T] motifs. Sites are only shown if present in one of the following representative strains: BG505 (clade A), AMC011 (clade B), CZA97 (clade C). The low conservation of the N332 site in clade AE is compensated by the high use of the N334 site (92%, AE; 21%, total).

| Data Collection Parameters | |
|---|---|
| Microscope | FEI Titan Krios |
| Voltage, kV | 300 |
| Detector | Gatan K2 Summit |
| Recording Mode | Counting |
| Magnification | 22,500x |
| Moive micrograph pixel size, Å | 1.31 |
| Dose rate, $e^-$/[(camera pixel)*s] | 10 |
| No. of frames per moive micrograph | 50 |
| Frame exposure time, ms | 200 |
| Movie micrograph exposure time, s | 10 |
| Total dose, $e^-/Å^2$ | 76 |
| Defocus range, µm | 1.5 to 3.0 |

| Map and Model Refinement Parameters | |
|---|---|
| No. of movie micrographs | 2184 |
| No. of molecular projection images in map | 68155 |
| Symmetry | C3 |
| Map resolution (FSC 0.143) | 3.79 |
| Map sharpening B-factor, $Å^2$ | -82.4 |
| No. of atoms in deposited model | 25821 |
| MolProbity score | 1.04 |
| C$\beta$ Outliers (%) | 0.00 |
| Rotamer Outliers (%) | 0.12 |
| Rama Outliers (%) | 0.71 |
| Clashscore | 0.95 |
| EMRinger score | 2.13 |
| EMDB | EMD-20224 |
| PDB ID | 6OZC |

**Table S1. CryoEM data collection and refinement parameters.**
